# Mild extraction preserves labile structural features of Gram-positive bacterial cell wall glycopolymers

**DOI:** 10.64898/2026.10.28.762426

**Authors:** Gordon Jacob Boehlich, Ahmad Tsjokajev, Pascal Mrozek, Mark Reglinski, Daniel Straume, Morten Kjos, Svein Jarle Horn, Helge C. Dorfmueller, Gustav Vaaje-Kolstad, Bjørge Westereng

## Abstract

Cell wall glycopolymers in Gram-positive bacteria are crucial for structural integrity, cell division, and stress resistance. Elucidating the structure of these glycopolymers is not only important for understanding fundamental bacterial physiology, but may facilitate vaccine development, phage therapy, antibiotic design and improvements in food fermentation safety. However, extraction protocols have so far relied on harsh chemicals that can damage these polymers, compromise their structural integrity, obscure their biological functions, and result in low yields. We present a straightforward method for isolating cell wall glycopolymers from Gram-positive bacteria that preserves essential structural features including attachment to peptidoglycan. The method was broadly applicable across phylogenetically diverse species and was not restricted to glycopolymer types. In total, we isolated 17 distinct glycopolymers from 11 species, including six with previously unreported structures, with a median yield of 49 mg per litre of culture. The simplicity and mild nature of the method, combined with the exceptional purity of the resulting products, has the potential to transform research on Gram-positive cell wall glycopolymers.

---

The cell envelope of Gram-positive bacteria is a complex, multilayered structure that plays crucial roles in cellular function and protection against hostile environments. In addition to peptidoglycan (PG), these bacteria produce cell wall glycopolymers (CWG) such as capsular polysaccharides (CPS)^1,2^, cell wall polysaccharides (CWPS)^3^, and anionic teichoic acids (TA)^4^ (**Figure 1**).

**Figure 1.**
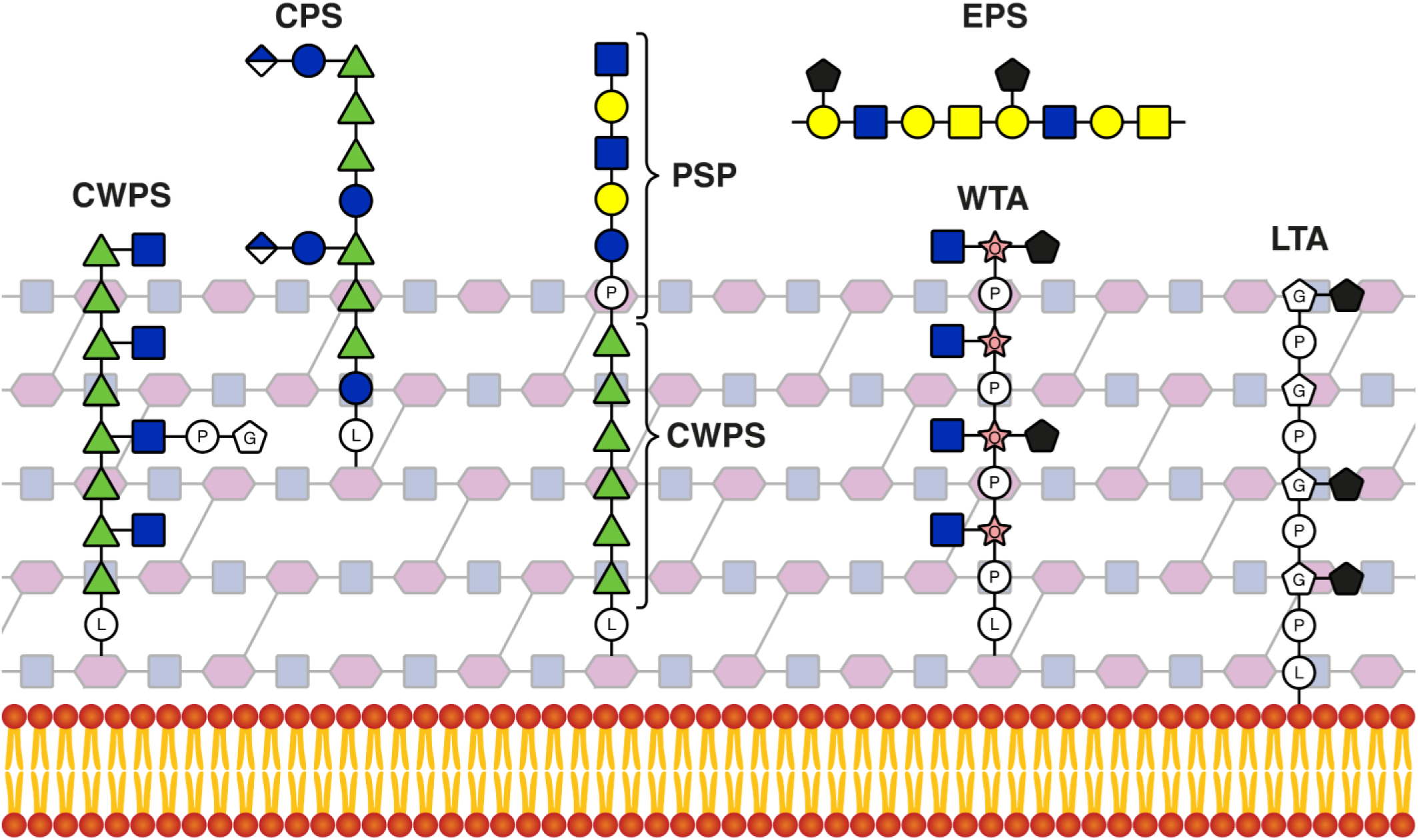
Examples of glycopolymers produced by Gram-positive bacteria. Translucent mesh of squares and hexagons: PG, repeating red circles with yellow tails: lipid bilayer, green triangle: Rha, blue square: GlcNAc, yellow square: GalNAc, blue circle: Glc, yellow circle: Gal, half-blue diamond: GlcA, pink star with O: ribitol, white pentagon with G: glycerol, black pentagon: non-carbohydrate modification, white circle with P: phosphate, white circle with L: linkage unit. The exact structure of the linkage unit and its attachment point to PG are known only for some bacteria^2,5,6^.

These CWGs, which can make up to 50% of the bacterial cell wall^7^, exhibit extensive structural diversity across bacterial taxa and among strains within individual species ^8–11^. The polymers fulfill a wide range of biological functions, including regulation of cell growth and morphology, tolerance to physicochemical stress, adhesion, colonization, pathogenicity, immune evasion, and resistance to antimicrobial agents^3,11^. Furthermore, their structures are also of interest for the development of antibiotics^12,13^ and vaccines^14–16^.

Structurally, TAs usually consist of repeating units of glycerol-phosphate (GroP) or ribitol-phosphate (RboP), which are often glycosylated or D-alanylated, contributing to their structural diversity^11,17^. TAs can be further divided into subgroups depending on their attachment mode to the bacterial cell wall. Lipoteichoic acids (LTA) are bound to the bacterial cell wall via a lipid anchor inserted into the lipid bilayer, while wall teichoic acids (WTA) are anchored covalently to the PG layer via a linkage unit^4^. Like WTAs, CWPS are usually covalently bound and embedded in the PG matrix, while CPS have higher weight-average molar mass (M_w_) and are located farther outside the PG layer^1,2^.

While some CPS have been reported to have covalent linkages to the cell wall, others may be associated by weaker interactions such as lectin binding, which can make them difficult to distinguish from exopolysaccharides (EPS), which are not bound to the cell wall^2,4,5^. Furthermore, hybrid structures consisting of an embedded CWPS and a surface-exposed polysaccharide pellicle (PSP) have also been reported^3,18–21^.

Extraction of CWGs that are covalently linked to the insoluble PG has usually relied on harsh chemical treatment with strong acids at elevated temperatures, toxic hydrofluoric acid that targets phosphodiester bonds, or strong bases^22–24^. These methods often compromise the structural integrity of the extracted CWG. For example, the presence of GroP in the Group A Carbohydrate (GAC) of *S. pyogenes*, which is of interest for vaccine development, remained unidentified for over 50 years due to the acid treatment cleaving phosphodiester bonds^25^.

To preserve the structural integrity of the CWGs, enzymatic degradation of the PG would therefore be preferred. This can be achieved by *N*-acetylmuramidases such as mutanolysin or lysozyme that cleave the β-1,4 bond between MurNAc and GlcNAc in the PG backbone. Previous studies on selected strains utilized mutanolysin to achieve this and purified the CWGs by chromatographic separation^25,26^. However, the high cost of mutanolysin, usually isolated from *Streptomyces globisporus*^27^, and the requirement for chromatographic separation limit the scalability of this approach. On the other hand, lysozyme is substantially less expensive because it can easily be isolated from hen egg white by ultrafiltration^28^.

The aim of this study was therefore to develop a method that (1) relies solely on affordable lysozyme for the release of CWGs while preserving their structural integrity; (2) avoids chromatographic purification steps whenever possible, thereby reducing time and resource requirements; and (3) is broadly applicable to a wide range of Gram-positive bacteria (**Figure 2**).

**Figure 2.**
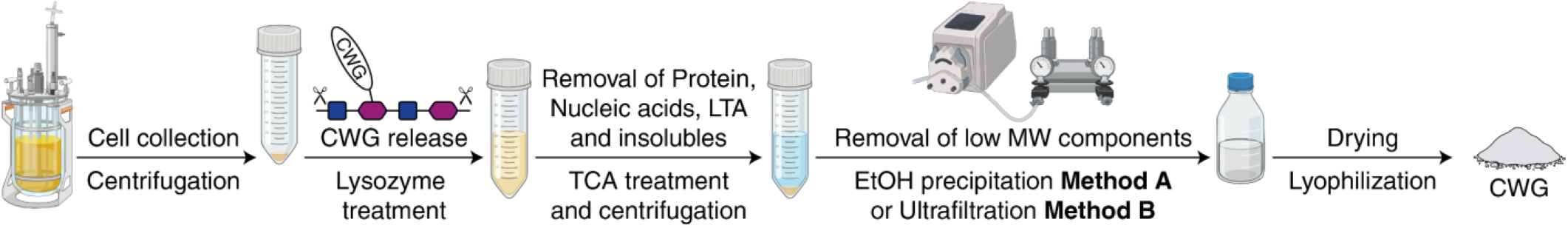
Procedures for CWG extraction involving lysozyme treatment. The method developed in this study utilizes lysozyme followed by a brief TCA-based cleanup and either ethanol precipitation (**Method A**) or ultrafiltration (**Method B**). Created in BioRender. Boehlich, G. J. (2026) https://BioRender.com/mio088a

## Results

### High loads of lysozyme are required to release CWG

To determine the amount of lysozyme required for the release of CWGs from the cell wall, *Lactococcus cremoris* TF121, which produces a sensitive CWPS-PSP copolymer, was used as a model organism to optimize the extraction method. Lysozyme (70,000–140,000 U/mg) amounts of 24, 60, 120, 240 and 480 mg per gram of wet cells were tested. As lysozyme remains active over a broad pH range (6-9)^29^, the reactions were performed without buffering to avoid introducing unnecessary buffer components, that will require removal during downstream processing. After the lysozyme extraction, CWG extracts underwent deproteinization for 1 h using TCA. Following deproteinization, the CWG was precipitated with ethanol and subsequently lyophilized (**Figure 2**, **Method A**).

Amounts of isolated CWG significantly increased with increasing amounts of lysozyme, plateauing at approximately 240 mg/g for wet cells of *L. cremoris* TF121 (**Extended Table 1**).

The use of mutanolysin was also investigated (**Extended Table 1, Entries 6 and 7**). Although the required enzyme dosage that maximized CWG yield was substantially lower than that of lysozyme, the cost of the enzyme required to achieve comparable results was 185-fold higher per mg of the pure polymer.

To determine purity, size, and structure the resulting materials were analyzed by size-exclusion chromatography (SEC) and nuclear magnetic resonance (NMR) spectroscopy (**Extended Figure 1**). SEC analysis using RI detection revealed that the elution profile of the CWG extracts became more defined with increasing lysozyme amounts. Similarly, ^1^H NMR spectra showed a substantial improvement in linewidth with increasing lysozyme load. The line broadening observed at lower lysozyme loads likely reflects slow molecular tumbling, indicating that substantial amounts of PG remained attached to the CWG. In the SEC-chromatograms, UV absorption at 280 nm was not observed at M_w_ >5000 kDa and minimal at M_w_ <5000 kDa across all conditions, suggesting negligible contamination by proteins or DNA. Furthermore, NMR spectra exclusively displayed signals attributable to bacterial cell-wall components, confirming that the materials were essentially free of contaminating nucleotides, proteins or lipids.

Only CWPS isolated using lysozyme loads of at least 240 mg/g of wet cells, as well as CWG extracted using mutanolysin, yielded NMR data suitable for precise structural characterization. Based on the gravimetric results, SEC data, and NMR data, a lysozyme dose of 240 mg/g of wet cells was used in all further experiments.

### Prolonged exposure to TCA may damage CWG

TCA is commonly used for purification of polysaccharide preparations. However, the low-pH environment may affect the structure of the extracted CWG. This was observed when CWPS from *L. cremoris* TF121, prior to polysaccharide precipitation with ethanol, was exposed to TCA for 24 h, resulting in a shift in the SEC profile toward lower M_w_ (**Extended Figure 2**). Additionally, peaks resembling oligosaccharides appeared at low M_w_.

The partial depolymerization may be explained by the presence of an anomeric glycosylphosphate and two galactofuranoses in the CWPS, which are highly susceptible to cleavage at pH 2 or lower, even at 4 °C. Limiting TCA exposure to 1 h preserved the polymer structure without affecting material purity. Therefore, exposure to TCA was limited to 1 h for all further experiments.

### Lysozyme-mediated extraction is applicable across phylogenetically diverse Gram-positive species

To evaluate the method across a phylogenetically diverse range of bacteria, the optimized protocol was applied to 14 Gram-positive strains representing 11 species, including multiple strains of several species. For strains whose CWG structures were known, ethanol precipitation was used to remove low M_w_ components to improve time efficiency (**Method A**). For strains with unknown CWG structures, ultrafiltration with a 5 kDa cut off membrane was used to fully ensure removal of all salts, as they may interfere with more complex NMR pulse sequences (**Method B**). It should be noted that yield and analytical data did not vary between the two methods for *L. cremoris* TF121 (**Table 1, entries 1 and 2**). Overall, CWGs were extracted from *L. cremoris*, *Lactococccus lactis* ssp. *lactis*, *Streptococcus pyogenes*, *Enterococcus faecalis*, *Streptococcus pneumoniae*, *Lactiplantibacillus plantarum, Lactiplantibacillus pentosus*, *Bacillus subtilis, Micrococcus luteus*, and *Staphylococcus aureus* with substantial yields of 1–36 mg per gram of wet cells. Total CWG amounts ranged from 9–1405 mg depending on the species, strain and culture volume which were more than sufficient for structural characterization by NMR in all cases (**Table 1**). SEC analysis of the CWG extracts showed multiple peaks in most samples. As these peaks were not baseline-separated, individual values of M_w_ and number-average molecular mass (M_n)_ could not be calculated. Nevertheless, the apparent peak molecular mass (M_p_) provides a close approximation of both M_w_ and M_n_ because the peaks appear to exhibit Poisson-like distributions whose dispersities (Ð) are near 1. The M_p_ indicated by these peaks were usually multiples of the lowest M_p_ peak, indicating that the CWGs were present as multimers of polymer chains. To validate the latter hypothesis, the different peaks observed for the various CWG extracts were separated by preparative SEC, collected as individual fractions, and analyzed by NMR (**Figure 3**).

**Figure 3.**
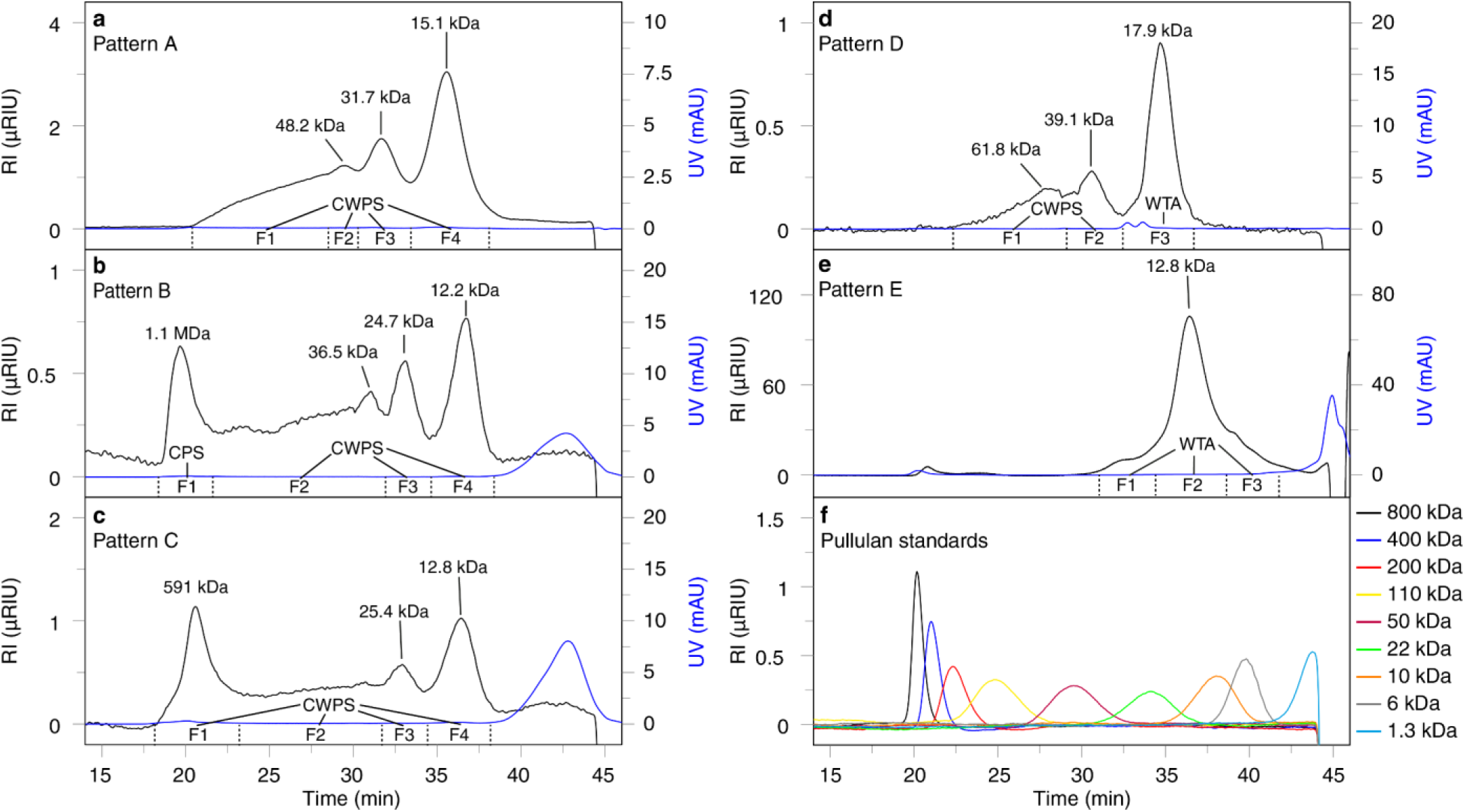
SEC analysis of CWGs from five different Gram-positive bacterial strains. Representative SEC profiles with refractive index detection (RI, black traces) and UV detection (mAU_280nm_, blue traces) of CWG isolates from five species and strains. (**a**) Multimers of a single CWG (**Pattern A,** *L. cremoris* TF121). (**b**) High M_p_ CPS and multimers of a CWG at lower M_p_ (**Pattern B**, *L. lactis* ssp. *lactis* CF131). (**c**) Multimers of a single CWG with an additional high M_p_ peak of the same CWG (**Pattern C**, *L. cremoris* TF117). (**d**) Two CWGs with similar M_p_ (**Pattern D**, *L. plantarum* WCFS1). (**e**) Nearly monomodal CWG (**Pattern E**, *S. aureus* Newman). (**f**) RI traces from pullulan standards of different molecular weights. Indicated apparent M_p_ are based on calibration with pullulan standards. Fractions used for NMR-based identification are indicated below the chromatogram by “F” followed by the corresponding fraction number.

**Table 1.**
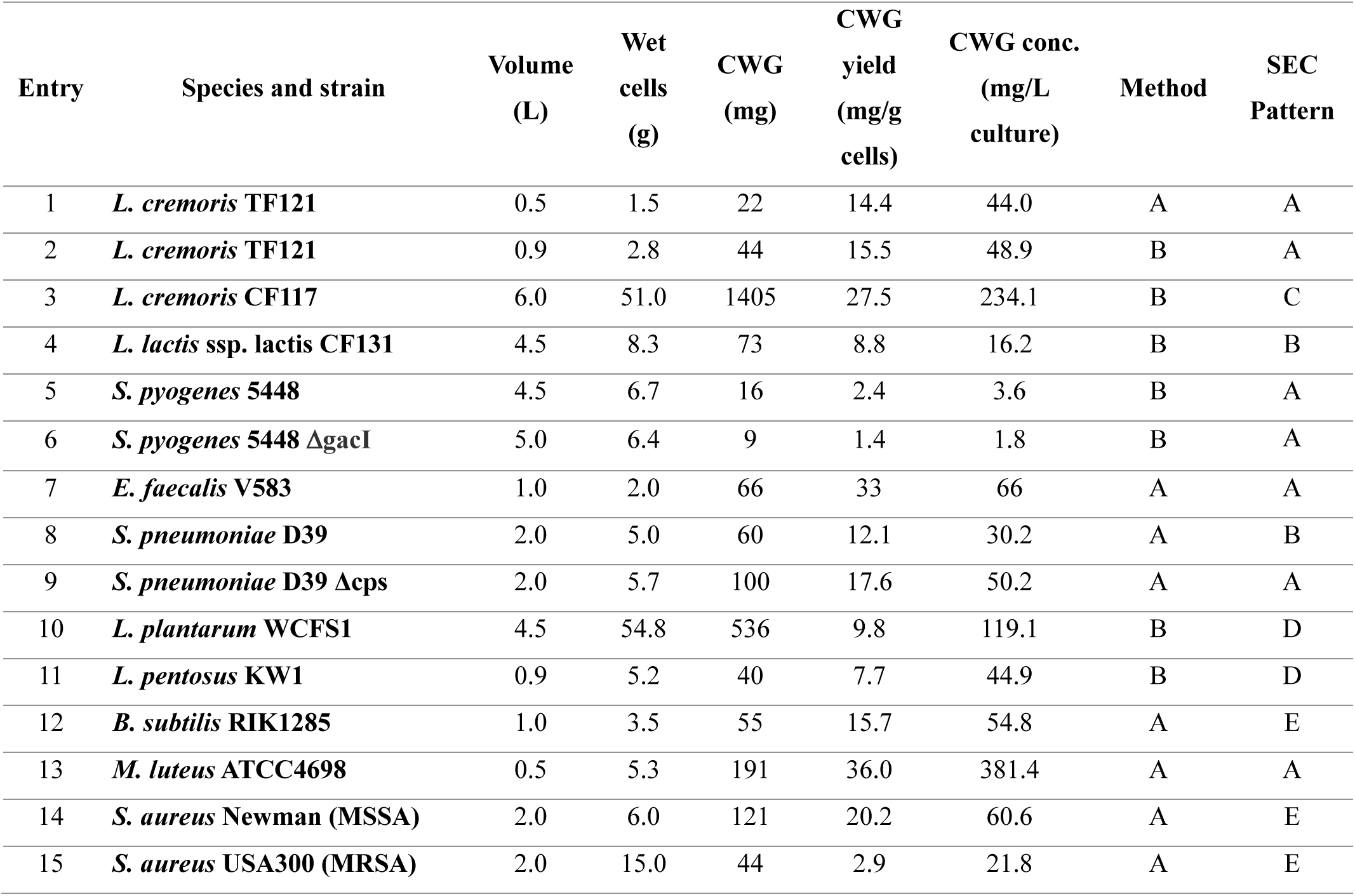
Cultivation volumes, cell amounts and gravimetric CWG yields for 15 extractions representing 14 bacterial strains. **Method A:** Isolation of CWG by EtOH precipitation. **Method B:** Isolation of CWG by ultrafiltration. SEC patterns A- E are depicted in Figure 3.

The resulting data confirmed the latter hypothesis for some of the bacteria, verifying that *L. cremoris* TF121, *S. pyogenes* 5448 and *M. luteus* ATCC4698 **only** produced a single CWG each (**Pattern A**). The SEC chromatograms of *L. lactis* ssp. *lactis* CF131 and *S. pneumoniae* D39 displayed additional peaks with high M_p_ which were determined to represent CPS (**Pattern B**). *L. cremoris* CF117 also displayed an additional high M_p_ peak, but NMR analysis revealed that it resulted from the same CWG as the low M_p_ peaks, which may be a result of less effective PG cleavage by the lysozyme for this strain (**Pattern C**). *L. plantarum* WCFS1 and *L. pentosus* KW1 both produced two different CWGs with similar M_p_ (**Pattern D**). Lastly, the CWG extracts from *B. subtilis* RIK1285 and both *S. aureus* strains showed mainly one peak, indicating that very few multimers of the CWG were present (**Pattern E**).

### Extracted CWGs display broad structural diversity

Having established that the extraction procedure yielded highly purified CWGs, we next examined their structural diversity. The obtained polymers were further characterized by NMR to determine their exact structures. For *L. cremoris* CF117 (Genotype A), the isolated CWPS featured the same rhamnan backbone as *L. lactis* UC509.9 (Genotype A)^31^, but the side chain was different: (1) the glucose substituent on the *O*-6 position of the GlcNAc residue was absent; (2) the terminal rhamnose was β-1-4-linked to the adjacent rhamnose, rather than α-1-4-linked; (3) the inner rhamnose residues were linked via an α-1-3 linkage instead of an α-1-2 linkage; (4) the galactose had no GroP modification (**Figure 4**).

**Figure 4.**
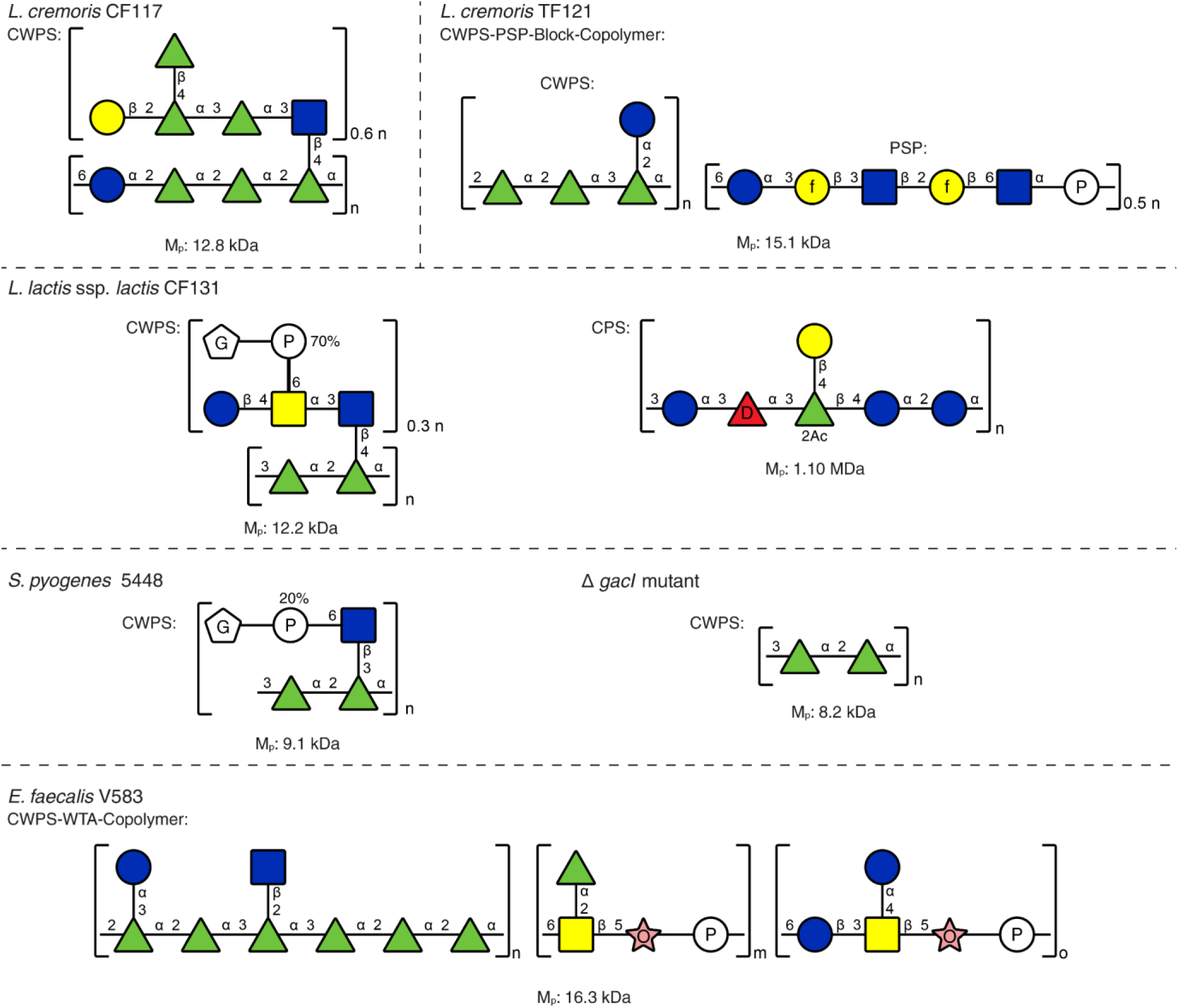
Structures and M_p_ of CWGs isolated from five rhamnan-CWPS-producing bacterial strains. Green triangle: L- Rha, red triangle with D: D-Fuc, blue square: D-GlcNAc, yellow square: D-GalNAc, blue circle: D-Glc, yellow circle: D-Gal, yellow circle with f: D-Galactofuranose, pink star with O: Rbo, white pentagon with G: Gro, white circle with P: phosphate. The reported M_p_ values are based on calibration with pullulan standards.

The CWPS of *Lactococcus cremoris* TF121 (Genotype C) showed NMR signals corresponding to two polysaccharides. These could not be separated by preparative SEC, suggesting that they are covalently linked, which is typical for genotype C lactococcal CWPS^19,21^. The copolymer consists of a rhamnan backbone, which is usually embedded further within the PG, and a polysaccharide pellicle (PSP) which is suggested to be presented outside the cell wall^18,20^. The rhamnan backbone matched the structure reported for *L. lactis* SK11 (Genotype C_3_)^21^, while the PSP structure matched that of *L. lactis* 3107^32^ (Genotype C_2_), making this a new combination (**Figure 4**).

The PSP contained two acid-labile galactofuranosides and an especially labile *N*-acetylglucosaminyl phosphate, all of which remained intact following the optimized extraction procedure. It has been shown that ^1^H-^13^C HSQC spectra of intact PSP can be obtained by means of solid-state MAS NMR of whole bacterial cells^32^. The liquid-state ^1^H-^13^C HSQC spectra of our extracted PSP matched perfectly with the reported solid-state data of the intact PSP, further supporting that the CWPS was not modified during extraction, which has not been achieved before for rhamnan-PSP copolymers from *Lactococcus lactis*.

Two CWGs were isolated from *Lactococcus lactis* ssp. *lactis* CF131. The high M_p_ polymer was elucidated as a new CPS composed of a hexasaccharide repeating unit consisting of D-glucose, D-galactose, L-rhamnose, and D-fucose in a molar ratio of 3:1:1:1. The rhamnose was fully acetylated at the *O*-2 position, indicating that no deacetylation occurred during the extraction (**Figure 4**). The structure of the low M_p_ rhamnan-CWPS isolated via lysozyme extraction from *L. lactis* ssp. *lactis* CF131 (Genotype B) was identical to that reported for *L. lactis* IL1403, as confirmed by matching NMR spectra (**Figure 4**)^33^.

This CWPS from *L. lactis* ssp. *lactis* CF131 and *L. lactis* IL1403 features the same alternating α-1,3 and α-1,2 rhamnan backbone as the Group A Carbohydrate (GAC) present in *S. pyogenes*, which, because of its surface exposure, antigenicity, and conservation, is of interest for vaccine development^14,15^. Applying the method to cells of *S. pyogenes* 5448 yielded the expected CWPS structure (**Figure 4**). The NMR data for the product were consistent with previously reported findings, including the partial glycerophosphate modification at the *O*-6 position of the GlcNAc, which had long remained unknown due to its lability^25^. Isolation of CWPS from an *S. pyogenes* 5448 mutant lacking the GlcNAc transferase gene, yielded a polysaccharide composed exclusively of the rhamnan backbone (**Figure 4**).^34^ Notably, extracts from both *S. pyogenes* strains contained poly-(GroP), which has previously been reported only as a component of *S. pyogenes* LTA. Previous studies have observed the release of LTA from *S. pyogenes* due to deacylation of the lipid anchor by enzymes present in the *S. pyogenes* membrane^35^. The resulting loss of the lipophilic substituents most likely prevented LTA removal by TCA precipitation, which successfully removed LTA from CWG extracts of all other strains.

Having demonstrated efficient recovery of GAC and its undecorated rhamnan core, we next examined whether the method could be applied to structurally distinct CWGs from other Gram-positive species. *Enterococcus faecalis* provided a particularly relevant example, as its enterococcal polysaccharide antigen (EPA) is a major cell-wall glycopolymer implicated in several important aspects of cell-envelope biology and pathogenesis. Applying the procedure to vancomycin-resistant *Enterococcus faecalis* V583 yielded a known rhamnan-WTA copolymer whose analytical data agreed with previous reports (Figure 4)^26^.

We next turned to *S. pneumoniae* D39, whose cell envelope contains multiple structurally distinct glycopolymers. In particular, the coexistence of a high-molecular-weight capsular polysaccharide and a structurally complex, phosphocholine-decorated WTA provided an opportunity to assess whether the procedure could recover and distinguish different CWGs from the same organism. CWG extraction of *S. pneumoniae* D39 cells yielded the expected high M_p_ CPS^36,37^ and phosphocholine-decorated low M_p_ WTA^38^. When the procedure was applied to cells of a mutant lacking the CPS biosynthesis gene, only the expected WTA was isolated (**Figure 5**).

**Figure 5.**
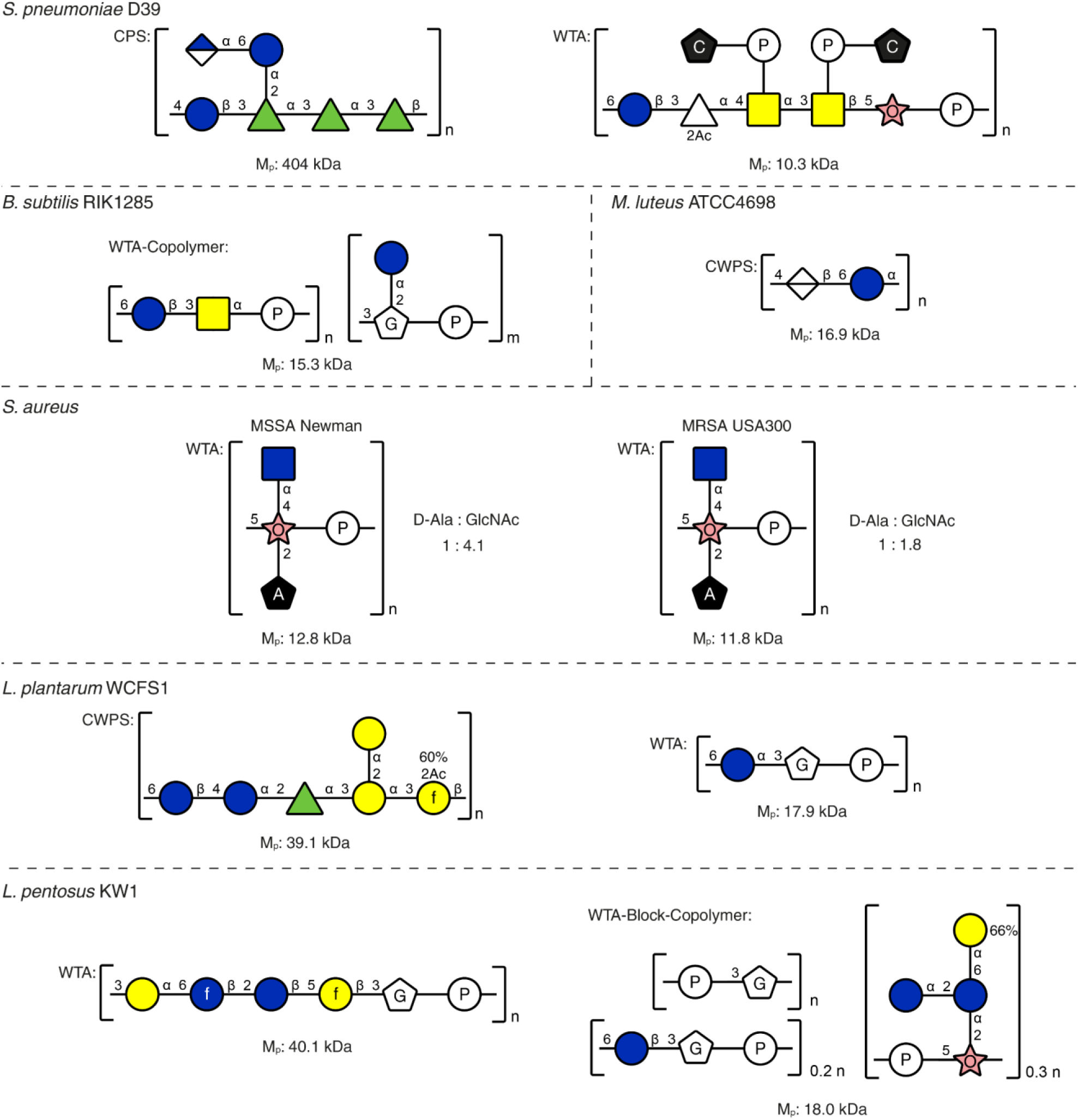
Structures and M_p_ of CWGs isolated from six teichoic and teichuronic acid-producing bacterial species. Green triangle: L-Rha; white triangle: D-4-N-FucNAc; blue square: D-GlcNAc; yellow square: D-GalNAc; blue circle: D-Glc; blue circle with f: D-Glucofuranose; yellow circle: D-Gal; yellow circle with f: D-Galactofuranose; half-blue diamond: D-GlcA; white diamond: D-ManNAcA; pink star with O: Rbo; white pentagon with G: Gro; white circle with P: phosphate; black pentagon with C: choline; D-alanine; black pentagon with A: D-alanine. The reported M_p_ values are based on calibration with pullulan standards.

We then investigated bacterial species commonly used in laboratory settings, which are known to produce various polyanionic CWGs. CWG extraction from cells of *Micrococcus luteus* ATCC4698, a strain commonly used in muramidase assays, yielded the expected teichuronic acid^39,40^. Furthermore, cells of *Bacillus subtilis* RIK1285, a bacterium commonly used for heterologous protein expression, yielded a WTA whose structure was identical to the WTA previously reported for *Bacillus subtilis* ssp. *subtilis* VKM B-501^T^ ^41^. Lastly, CWG extraction from *S.aureus* MSSA Newman and MRSA USA300 cells yielded the expected RboP-based WTA, decorated with α-GlcNAc and D-alanine esters^42^. In comparison to the methicillin-sensitive Newman strain, the methicillin-resistant USA300 strain displayed a higher degree of D-alanylation (**Figure 5**).

Furthermore, we investigated whether the procedure could reveal previously uncharacterized CWG diversity in WTA producing species. For this purpose, we examined *L. plantarum* WCFS1, a well-characterized model lactic acid bacterium with an established WTA structure. Two CWGs were extracted from the cell pellet of *L. plantarum* WCFS1. One was the previously reported WTA, composed of a simple α-glucosyl-glycerophosphate repeating unit^43^, while the other was a new CWPS composed of a hexasaccharide repeating unit consisting of rhamnose, glucose, and galactose in a molar ratio of 1:2:3. Notably, one of the galactosides in this polysaccharide is a labile furanoside that is partially *O*-acetylated at the *O*-2-position (**Figure 5**).

Lastly, for *L. pentosus* KW1^44^, which is closely related to *L. plantarum,* the procedure yielded two polymers which were identified as new teichoic acids. The teichoic acid of higher M_p_ had a strict repeating unit composed of glucose, galactose and GroP in a molar ratio of 2:2:1. Notable labile features of the structure were the glucofuranoside, the galactofuranoside, and the phosphate linkage, all of which were preserved after lysozyme extraction (**Figure 5**). The other teichoic acid was a block polymer consisting of three different subunits linked by phosphodiester bonds. The most abundant subunit by numerical amount was identified as GroP. The next most prevalent subunit was identified as RboP bearing a kojibiose substituent at the 2-position. Roughly two-thirds of this kojibiose carried an additional galactose substituent. The final subunit was identified as β-glucosyl-GroP (**Figure 5**).

### CWG isolates contain residual PG fragments

Since WTA and CWPS are covalently linked to PG, fragments from PG degradation remain attached and can be observed in NMR spectra at lower intensities. PG structures display considerable structural variation in the stem peptide, both in amino acid composition and in the mode of crosslinking^45^. Characteristic signals stemming from these PG fragments can be observed between 1 and 3 ppm in ^1^H NMR spectra. Two features are especially useful when identifying these signals as PG fragments, as they resonate at frequencies not occupied by carbohydrate signals: (1) the broad triplet at ∼2.40 ppm arising from CH_2_ groups in γ-glutamate or isoglutamine; (2) multiple overlapping doublets between 1.36 and 1.40 ppm stemming from CH_3_ groups of alanine and lactate (**Extended Figure 3**). Notably, signals of CH_3_ groups did not overlap with the CH_3_ signals of rhamnoses between 1.25 and 1.35 ppm or D- alanyl esters at 1.64 ppm on a 9.4 T (“400 MHz”) spectrometer.

The PG fragment signals can be utilized to gain further insight into the PG structure of the respective strain. The greatest variation between structures is the diamino acid, which is essential for crosslinking PG via the interpeptide bridge. ^1^H-^13^C HSQC showed that the stem peptide in *L. plantarum* WCFS1 contained *meso*-diaminopimelic acid (*m*-DAP) as the diamino acid, whereas *L. cremoris* TF121 displayed signals typical for lysine (**Extended Figure 3**). Furthermore, signals corresponding to asparagine, a characteristic feature of lactococcal interpeptide bridges, were observed in the CWPS of *L. cremoris* TF121. Likewise, signals corresponding to the typical pentaglycine crosslinks present in *S. aureus* were observed (**Extended Figure 3**). Most extracts also showed small signals from *O*-acetyl groups, which were not associated with the CWPS. These signals most likely originated from *O*-acetyl groups in the PG residues, which have been linked to lysozyme resistance^46^.

NMR data can also give insight into the crosslinking of PG (**Extended Figure 4**). The CWPS of *L. cremoris* TF121 displayed a signal at 3.20 ppm for the CH_2_ group next to the primary amine in the ^1^H NMR spectrum, which was broadened because it was crosslinked to another peptide subunit. This signal was also present in the CWPS of *L. cremoris* CF117. However, an additional triplet also stemming from a lysine was present at 3.00 ppm. The narrower lineshape of this triplet indicates that this signal stems from non-crosslinked lysine. This hypothesis was confirmed by HMBC, where the broadened signal at 3.20 ppm showed a correlation to asparagine, unlike the sharp triplet at 3.00 ppm. These results indicate that the PG in *L. cremoris* CF117 is considerably less crosslinked than the PG in *L. cremoris* TF121.

The PG fragments present in the CWPS extract of *L. cremoris* CF117 could be degraded by alkaline treatment at elevated temperatures (**Extended Figure 5**). After this treatment and removal of low M_w_ components by ultrafiltration, the ^1^H NMR of the product showed mainly CWPS signals, while the PG signals were mostly not present. Furthermore, the multimodal size distribution observed in the SEC chromatogram collapsed into a monomodal distribution, indicating that the multimodal distributions observed in SEC-chromatograms (**Figure 3**) were in fact caused by multiple CWPS polymer chains attached to the same PG residues.

## Discussion

The method developed in this work enables rapid and mild extraction of structurally intact CWGs from Gram-positive bacteria, which is scalable due to the affordability of hen egg white lysozyme and minimal downstream processing. While the required concentration of lysozyme may seem high, it should be noted that its concentration in hen egg whites is between 2.2 and 4.5 mg/mL^47^. This concentration is only 5- to 10-fold lower than the concentration we determined to be necessary for near-complete degradation of the PG from a large amount of bacterial cells. The method successfully extracted CWGs from all strains of the eleven species tested in this study, yielding 17 distinct CWGs. Notably, the high concentrations of lysozyme were able to successfully release WTA from cells of *S. aureus*, which has been deemed lysozyme resistant^46^, without requiring PG *O*-deacetylation or addition of lysostaphin for the degradation of the pentaglycine crosslinks. Furthermore, TCA proved to be surprisingly effective in removing non- CWG components and did not affect the CWG structure as long as the exposure time was kept to one hour. Degradation of protein and DNA, commonly done by addition of proteinases or DNases, was therefore not needed for any samples in the present study.

Following the TCA precipitation, both ethanol precipitation (**Method A**) and ultrafiltration (**Method B**) provided effective routes for CWG recovery. While EtOH precipitation of CWP (**Method A**) was convenient for salt removal in small scale extracts and provided material suitable for verification by NMR analysis, ultrafiltration (**Method B**) appeared to be a better alternative for large scale extractions where the use of large volumes of ethanol would be impractical. Further chromatographic purification was only necessary for species or strains producing multiple CWGs, eliminating another time- and resource-consuming step compared to conventional methods.

The yields of CWGs were strain-specific, ranging from ca 2-400 mg/L of culture. The lower end of yields was observed for *S. pyogenes*, which did not grow to high cell densities due to autolysis. The yield from *Micrococcus luteus* was the highest, which is consistent with its well-known sensitivity to enzymatic PG cleavage but could also be caused by higher CWPS production. Overall, the median yield of 49 mg per liter of culture suggests that one liter of bacterial culture is more than sufficient for isolating CWG amounts suitable for characterization and further experiments in most cases. Further improvements could be achieved by optimizing growth conditions for individual strains, although such optimization may be warranted only when large quantities of a particular CWG are required.

Because the method relies solely on PG degradation, the extracted CWGs retain covalently linked PG fragments, which also causes the appearance of apparent multimers in SEC. The residual PG fragments can provide insight into PG structure, including peptide-side-chain composition and the degree of cross-linking. Due to the insolubility of PG, such data are usually obtained from solid-state NMR or LC-MS after enzymatic solubilization. The high resolution of liquid-state NMR can allow detection of modifications and precise determination of linkages previously missed by these techniques. It should be noted, however, that the PG fragments are only representative of the PG structure around the CWG attachment loci. Furthermore, structural information about the linkage between CWGs and PG may also be gained from these extracts. However, such an analysis would require either highly sensitive instrumentation or an enzyme capable of depolymerizing the respective CWG, because the residual PG fraction is relatively small. Structural identification of the linkages between CWGs and PG may also aid in identifying enzymes to remove the PG residues from the CWG. In the case of *L. cremoris* CF117 CWPS, PG removal could be achieved under strongly alkaline conditions. However, such treatment is unsuitable for most CWG structures because it promotes cleavage of phosphate and carboxylate esters and can also result in partial loss of *N*-acetyl groups.

The method described in the present study may also provide a means to differentiate between CWPS, CPS and EPS. Some polysaccharides are referred to as EPS solely because they have been recovered from the supernatant. Cell lysis can also release CPS and CWPS into the supernatant in some cases, leading to misclassification as EPS. The term EPS usually refers to secreted polysaccharides that are not covalently linked to the bacterial cell wall.^6^ For example, *L. pentosus* KW1^48^ has been reported to release a high-M_w_ EPS into the culture supernatant; this polymer was not recovered from the cells by our method, consistent with it not being covalently attached to the cell wall. For smaller CWG (<50 kDa), the presence of residual PG signals in NMR is also a strong indication of covalent linkage to the cell wall.

In conclusion, our simple and mild workup method allows for the isolation of CPS, WTA and CWPS containing labile features such as furanosides, acylations, and phosphate linkages, which are often not detected using conventional methods. Consequently, we were able to identify and characterize six unpublished structures. This broadly applicable procedure has the potential to transform the study of Gram-positive cell wall glycopolymers by enabling detailed characterization of their structures, their covalent connections to PG, and ultimately their structure– function relationships.

## Acknowledgements

The authors thank Dr. Philipp Garbers, Dr. Kamilla Wiull, Prof. Geir Mathiesen, and Assoc. Prof. Davide Porcellato for kindly providing bacterial stocks. Furthermore, we thank Synnøve Sætervik and Cornelia Rognstad Karlsen for invaluable help with isolation of CWGs and Zhian Salehian for bacterial cultivation.

## Funding

This study was supported by Research Council of Norway project 326272 and internal strategic funding provided by The Norwegian University of Life Sciences. Laboratory facilities used in this study were financed through RCN infrastructure grants 270038, 295910 and 296083. MR and HD are funded by the RIGHT Foundation RF-2023-V2 and Wellcome 225350/Z/22/Z.

## Declaration of Competing Interests

Gordon Jacob Boehlich, Ahmad Tsjokajev, Pascal Mrozek, Gustav Vaaje-Kolstad and Bjørge Westereng have submitted a patent application covering the CWG extraction method described in this study.

## Experimental

### Bacterial cultivation and fermentation procedures

#### Bacterial strains

In-house strains, L. pentosus KW1, L. cremoris CF117, L. lactis ssp. lactis CF131, L. cremoris TF121, S. pyogenes 5448 ΔgacI mutant, and publicly available strains L. plantarum WCFS1, S. pyogenes 5448, E. faecalis V583, S. pneumoniae D39, M. luteus ATCC4698, B. subtilis RIK1285, S. aureus Newman and S. aureus USA300 were cultivated for CWG extraction.

#### Fermentation of Lactiplantibacilli and Lactococci

Fermentations of *L. pentosus* KW1 and *L. plantarum* WCFS1 were conducted using a modified version of De Man, Rogosa, and Sharpe (MRS) medium. The medium contained 10 g/L peptone from casein, 8 g/L meat extract, 4 g/L yeast extract, 2 g/L K_2_HPO_4_, 5 g/L sodium acetate trihydrate, 2 g/L ammonium citrate tribasic, 0.2 g/L MgSO_4_·7 H_2_O and 0.05 g/L MnSO_4_·H_2_O. Fermentations of *L. lactis* strains were performed in M17 broth with glucose instead of lactose (0.5 g/L ascorbic acid, 0.25 g/L MgSO_4_, 5 g/L meat extract, 2.5 g/L meat peptone, 19 g/L sodium glycerophosphate, 5 g/L soy peptone (papainic), 2.5 g/L yeast extract).

High M_w_ polysaccharides and polymers were removed from the MRS and M17 media by ultrafiltration prior to bacterial cultivation.^49^ Ingredients equivalent to 5 L of MRS and M17 medium were separately dissolved in 1 L of Milli-Q water and ultrafiltered using a Pellicon® 2 Mini Cassette Holder (Millipore) equipped with a Biomax® 30 kDa membrane (P2B030V01, Millipore), with a peristaltic pump used for circulation, until the permeate volume reached 5 L. The resulting permeates were sterile-filtered using 0.22 µm pore size Steritop filters (SCGPS05RE, Millipore) and used as fermentation media. Prior to inoculation, the medium was adjusted to pH 6.0 using 2.5 M H_2_SO_4_ and supplemented with D-glucose to a final concentration of 20 g/L.

*Lactiplantibacilli* and *Lactococci* were fermented in 1.5 L Minifors 2 bioreactors (Infors HT, Switzerland). *Lactiplantibacillus* strains were cultivated in ultrafiltered MRS medium at 37 °C with stirring at 200 rpm. The pH was maintained at 6.0 by automatic addition of 5 M NaOH. *Lactococcus* strains were cultivated in ultrafiltered M17 medium under identical conditions, except at a temperature of 18 °C. The inocula for both genera were prepared by inoculating 40 mL of growth medium in a 50 mL Nunc plastic tube with bacterial cells from a -80 °C glycerol stock using an inoculation loop, followed by overnight incubation at 37 °C and 18 °C, respectively.

#### Cultivation of *S. pyogenes*

The *S. pyogenes* strains were cultivated in Todd-Hewitt broth (THB) supplemented with yeast extract (THY). The THY medium contained 3.1 g/L beef heart solids from infusion, 20 g/L peptic digest of animal tissue, 2 g/L D- glucose, 2 g/L NaCl, 0.4 g/L Na_3_PO_4_, 2.5 g/L Na_2_CO_3_ and 5 g/L yeast extract.

Cultivation of *S. pyogenes* strains was carried out in 1 L Duran bottles using THY medium, incubated overnight at 37 °C under 5 % CO_2_. After cultivation, the cultures were autoclaved at 121 °C for 22 min for safety reasons. The inoculum was prepared by inoculating 50 mL of THY medium with 2–3 colonies of each strain, followed by incubation for 7 h at 37 °C and 5 % CO_2_.

#### Cultivation of *E. faecalis*

Cultivation of *Enterococcus faecalis* V583 was carried out in 2 L baffled flasks containing 500 mL double yeast tryptone (dYT) medium (16 g/L tryptone, 10 g/L yeast extract, 5 g/L NaCl), incubated overnight at 37 °C with 200 rpm shaking. The inoculum was prepared by inoculating 5 mL dYT medium from a cryostock, followed by incubation overnight at 37 °C with shaking at 200 rpm.

#### Cultivation of *S. pneumoniae*

Cultivation of *S. pneumoniae* strains was carried out by inoculating 2 L THB (20 g/L peptone, 3.1 g/L beef heart solids from infusion, 2.5 g/L Na_2_CO_3_, 2 g/L glucose, 2 g/L NaCl, 0.4 g/L Na_2_HPO_4_) to an OD_550_ of 0.05. Cells were grown at 37 °C until reaching an OD_550_ of 0.5.

#### Cultivation of *M. luteus*

Cultivation of *Micrococcus luteus* ATCC 4698 was carried out in 2 L baffled flasks containing 500 mL tryptic soy broth (TSB) medium (17 g/L pancreatic digest of casein, 3 g/L peptic digest of soybean, 5 g/L NaCl, 2.5 g/L K_2_HPO_4_, 2.5 g/L glucose), incubated for 72 h at 30 °C with 200 rpm shaking. The inoculum was prepared by inoculating 5 mL TSB medium from a cryostock, followed by incubation for 48 h at 30 °C with 200 rpm shaking.

#### Cultivation of *B. subtilis*

Cultivation of *Bacillus subtilis* RIK1285 was carried out in 2 L baffled flasks containing 500 mL lysogeny broth (LB) medium supplemented with 0.2% glucose (10 g/L tryptone, 5 g/L yeast extract, 10 g/L NaCl, 2 g/L glucose), incubated overnight at 37 °C with 200 rpm shaking. The inoculum was prepared by inoculating 5 mL LB medium supplemented with 0.2% Glucose from a cryostock, followed by incubation overnight at 37 °C with shaking at 200 rpm.

#### Cultivation of *S. aureus*

Cultivation of *S. aureus* strains was carried out in 3 L Erlenmeyer flasks containing 1 L brain heart infusion (BHI) medium (200 g/L infusion from calf brains, 250 g/L infusion from beef hearts, 10 g/L proteose peptone, 2 g/L D- glucose, 5 g/L NaCl, 2.5 g/L Na_2_HPO_4_) incubated overnight at 37 °C with shaking at 200 rpm. Cultures were inoculated 1:1000 from a preculture in BHI.

#### Extraction, isolation, and purification of cell wall glycopolymers from cells

The cell suspensions from growth cultures were centrifuged (JLA-8.1000, Avanti JXN-26, Beckman Coulter) at 15,970 × g and 4 °C for 30 minutes. The supernatant was discarded and CWGs from the cell pellet were obtained as shown in Figure 2. The pellet was washed once with Milli-Q water and weighed. For each gram of wet cells, 12 mL of lysozyme solution (20 mg/mL; 70 000–140000 U/mg, Sigma-Aldrich, Merck, Darmstadt, Germany) was added and the mixture incubated overnight at 37 °C with stirring at 150 rpm. TCA (50 %, w/v) was then added to the reaction mixture to a final concentration of 5 %, followed by incubation at 4 °C for 30 minutes. The mixture was centrifuged using a swing-bucket rotor (Megastar 1.6R, VWR, Germany) at 4136 × g for 30 minutes at 4 °C and the supernatant immediately neutralized to pH 7 with 1 M NaOH and filtered using 0.22 µm pore size Steritop filter. The CWG in the filtrate was then isolated by one of two methods:

**Method A**: The CWG was precipitated by adjusting the filtrate to 80% EtOH using 96% EtOH and incubating the mixture at 4 °C for 16 h. The CWG was then collected by centrifugation using a swing-bucket rotor (Megastar 1.6R, VWR, Germany) at 4136 × g for 10 minutes at 4 °C. After discarding the supernatant, the CWG was then redissolved in deionized water and lyophilized to yield the CWG as a colorless solid.

**Method B:** The CWG was purified by ultrafiltration and diafiltration of the filtrate using a Biomax® 5 kDa membrane (P2B005V01, Millipore). The retentate was then collected and lyophilized to yield the CWG as a colorless solid.

### Characterization of cell wall glycopolymers

#### Size-exclusion chromatography

The SEC setup included a Yarra SEC-4000 column and a Yarra SEC-2000 column (300 × 7.8 mm, 3 µm), both from Phenomenex (Torrance, CA, USA). The columns were connected in series to a Dionex™ UltiMate™ 3000 HPLC system (Thermo Fisher Scientific, Waltham, MA, USA). Elution was performed at a flow rate of 0.5 mL/min using 0.1 M Na_2_SO_4_ as the mobile phase. Detection was carried out using UV absorbance at 280 nm (Thermo Scientific Dionex VWD-3000 or VWD-3400 detector) and refractive index (ERC RefractoMax 520, Malvern Viscotek TDA 305 or Thermo Scientific Vanquish VC-D60-A01). The column setup had an exclusion limit of 1500 kDa, and injection volumes were 20 µL. Pullulan standards (1.3–800 kDa; Agilent Technologies, Mainz, Germany) were regularly run to monitor column performance and calculate apparent peak polymer masses (M_p_). M_p_ were calculated by conventional calibration^50^ using **Equation 1**.

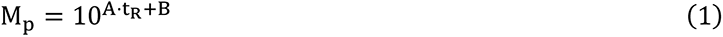

M_p_ = peak molecular mass in Da

t_R_ = retention time at peak maximum in min

A = -0.2802 for t_R_ < 22.3 min and -0.083 for t_R_ ≥ 22.3 min

B = 11.533 for t_R_ < 22.3 min and 7.1316 for t_R_ ≥ 22.3 min

#### Preparative Size-exclusion chromatography

To determine the identity of different peaks in analytical SEC chromatograms, preparative SEC was employed for fractionation. This was conducted using an Agilent 1260 Infinity Preparative HPLC system equipped with Yarra SEC- 3000 PREP and SEC-2000 PREP columns (300 × 21.2 mm, 5 µm; Phenomenex), connected in series. Detection was achieved via refractive index and dual-wavelength UV at 210 nm and 280 nm. The mobile phase consisted of 0.08 M ammonium formate (NH₄^+^HCOO^-^), pH 4.0, with a flow rate of 5 mL/min at room temperature. The injection volumes were 4.5 mL, corresponding to 25 mg of dry sample per injection. Collected fractions were lyophilized to remove water and ammonium formate.

#### NMR analysis

All materials analyzed by NMR were dissolved in D_2_O and subsequently lyophilized to minimize the HDO signal. Finally, samples were dissolved in 0.45 mL D_2_O containing 0.05 wt. % 3-(trimethylsilyl)propionic-2,2,3,3-*d*_4_ acid, sodium salt. The samples were measured using 5 mm Norell® Sample Vault Series™ NMR tubes. Spectra were recorded at 333 K unless otherwise noted using a Bruker AVANCE III HD 400 MHz instrument (Bruker BioSpin, Ettlingen, Germany) equipped with a BBO room-temperature probe. Chemical shifts are reported in ppm with Na- Trimethylsilylpropanoate-*d*_4_ (δ ^1^H/^13^C = 0.00 ppm) as internal standard. ^31^P shifts are reported using 85% phosphoric acid in D_2_O as external standard (δ ^31^P = 0.00 ppm). Coupling constants (*J*) are reported in Hz. All materials were analyzed by 1D ^1^H and 2D ^1^H-^13^C HSQC NMR spectroscopy. If the data for a CWG did not match those of a previously reported structure, the structure was further elucidated by NMR.^51,52^ In short: monosaccharides were identified by TOCSY and HSQC-TOCSY. The signals were then assigned to the corresponding ring positions using COSY, RELAY-COSY and H2BC. Linkages were then identified by HMBC. If the linkages could not be identified by HMBC, NOESY spectra were used for identification. Anomeric configurations were determined by ^1^*J*_CH_ coupling constants and NOESY. ^13^C shift values were taken from DEPTQ spectra and ^31^P shift values from ^1^H-decoupled ^31^P- spectra. The following pulse programs were used: “zg30” for ^1^H, “deptqgpsp.2” for DEPTQ-^13^C, “zgpg30” for ^31^P, “cosygpmfppqf” for ^1^H-^1^H COSY, “cosyqfrl” for RELAY-COSY, “cosyqfr2” for double RELAY-COSY, “dipsi2gpphzs” for ^1^H-^1^H-TOCSY, “hsqcetgpsisp2.2” for HSQC, “h2bcetgpl3” for H2BC, “hsqcdietgpsisp.2” for HSQC-TOCSY, “hmbcetgpl3nd” for HMBC, and “noesygpphpp” for NOESY. If resolution in the ^13^C dimension was not sufficient, 2D-selective versions of HSQC, HMBC and HSQC-TOCSY spectra were acquired.

#### Monosaccharide composition analysis of new CWG structures

To further confirm the NMR-based identification of monosaccharides in newly characterized CWGs, high-performance anion-exchange chromatography with pulsed amperometric detection (HPAEC-PAD) was performed. To convert the polysaccharides to monosaccharides, a 1 mg/mL solution of the CWG sample was hydrolyzed in 1 M trifluoroacetic acid (TFA) at 100 °C for 1 h. The hydrolysate was dried in a Concentrator plus (Eppendorf, Hamburg, Germany) and redissolved in Milli-Q water to a final concentration of 1 mg/mL. The sample was further diluted to 0.5 and 0.1 mg/mL with Milli-Q water and analyzed using a Dionex™ ICS-6000 System (Thermo Fisher Scientific) equipped with a Dionex™ CarboPac™ PA1 column (2 × 250 mm). The mobile phase consisted of 12 mM KOH with a flow rate of 0.6 mL/min at 30 °C. An injection volume of 0.4 µL was used. Analytical standards, including glucose (Glc), galactose (Gal), glucosamine (GlcN), galactosamine (GalN), rhamnose (Rha), fucose (Fuc) and mannose (Man), were purchased from Sigma-Aldrich (Merck, Darmstadt, Germany) and prepared identically to the CWG samples.

#### Determination of monosaccharide absolute configuration in new CWG structures

20 mg of dry CWG powder was hydrolyzed in 2 mL of 1 M HCl at 100 °C for 1 h to accomplish complete depolymerization. The hydrolysate was then neutralized with solid Ag_2_CO_3_ and 100 µL of 100% Ac_2_O was added to reacetylate the amino sugars. The mixture was stirred for 1 h at room temperature, followed by filtration through a 0.45 µm pore size PES syringe filter (Sarstedt, Filtropur S). Dowex 50W X8 in hydrogen form was added and the mixture was immediately filtered again using the same filter type. The water was removed in vacuo and the absolute configurations of monosaccharides were determined by reversed-phase chromatography (RP-HPLC) using a COSMOSIL 5C18-AR-II column (4.6 × 250 mm, 5 µm, 120 Å; Phenomenex, Torrance, CA, USA) after derivatization with L-cysteine methyl ester and *o*-tolyl isothiocyanate.

#### Removal of PG residues from *L. cremoris* CF117 CWPS

100 mg of *L. cremoris* CF117 CWPS extract was dissolved in 10 mL of 1M NaOH and heated at 100 °C for 1 h. The mixture was purified by ultrafiltration and diafiltration of the filtrate using a Biomax® 5kDa membrane (P2B005V01, Millipore). The retentate was then collected and lyophilized to yield 67 mg PG-free CWPS as a colorless solid.

---

**Extended Figure 1.**
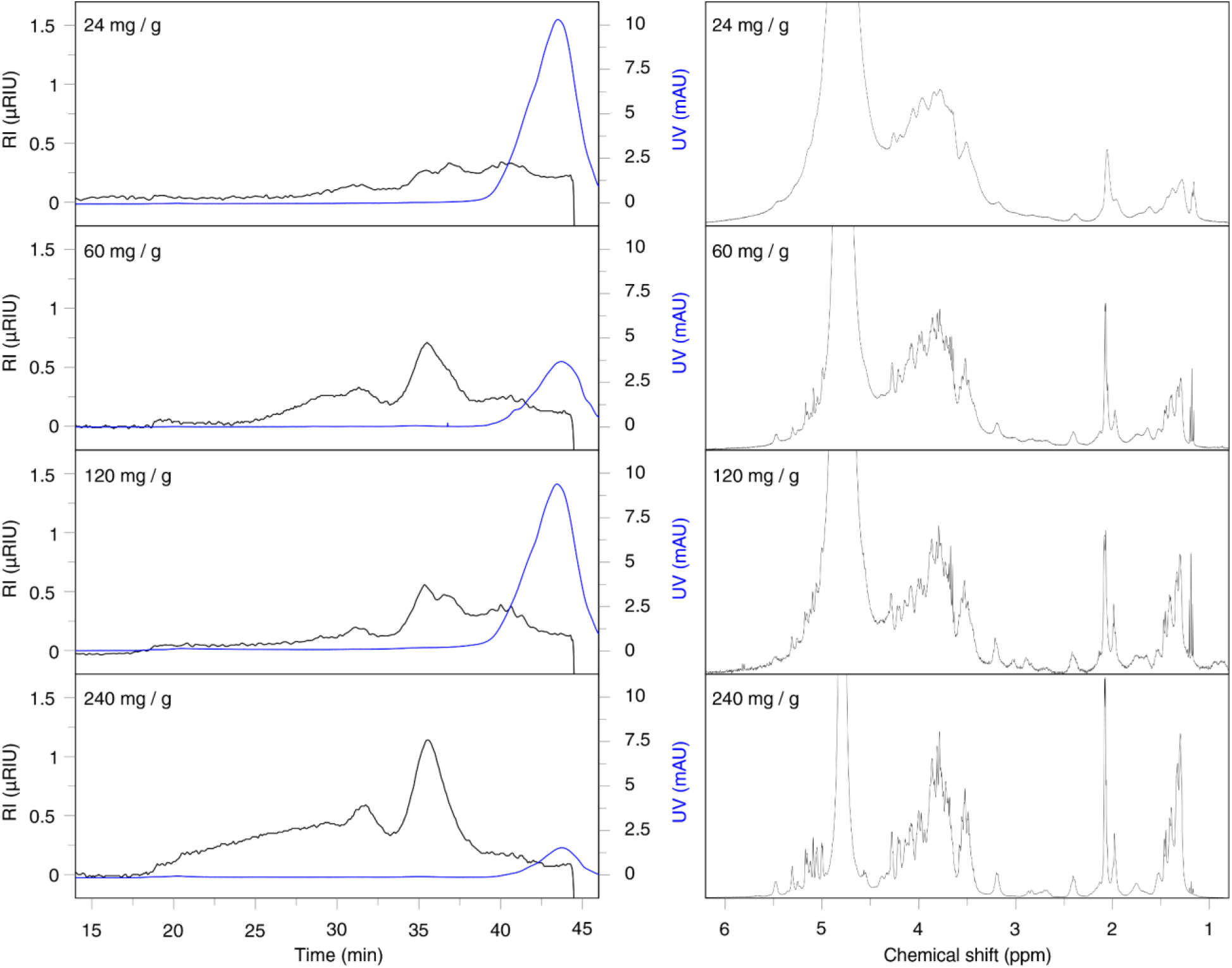
Analysis of CWG by SEC and NMR at different lysozyme loads. Left: Size-exclusion chromatography (SEC) with refractive index detection (RI, black traces) and UV detection (mAU_280nm_, blue traces) of *L. cremoris* TF121 CWG isolates obtained using increasing lysozyme loads. **Right:** ^1^H NMR spectra (400 MHz, D_2_O, 298 K) of CWG isolates corresponding to the same lysozyme loads. The lysozyme load per gram wet cells used for extraction is indicated in the upper left corner of the chromatograms and spectra.

**Extended Figure 2.**
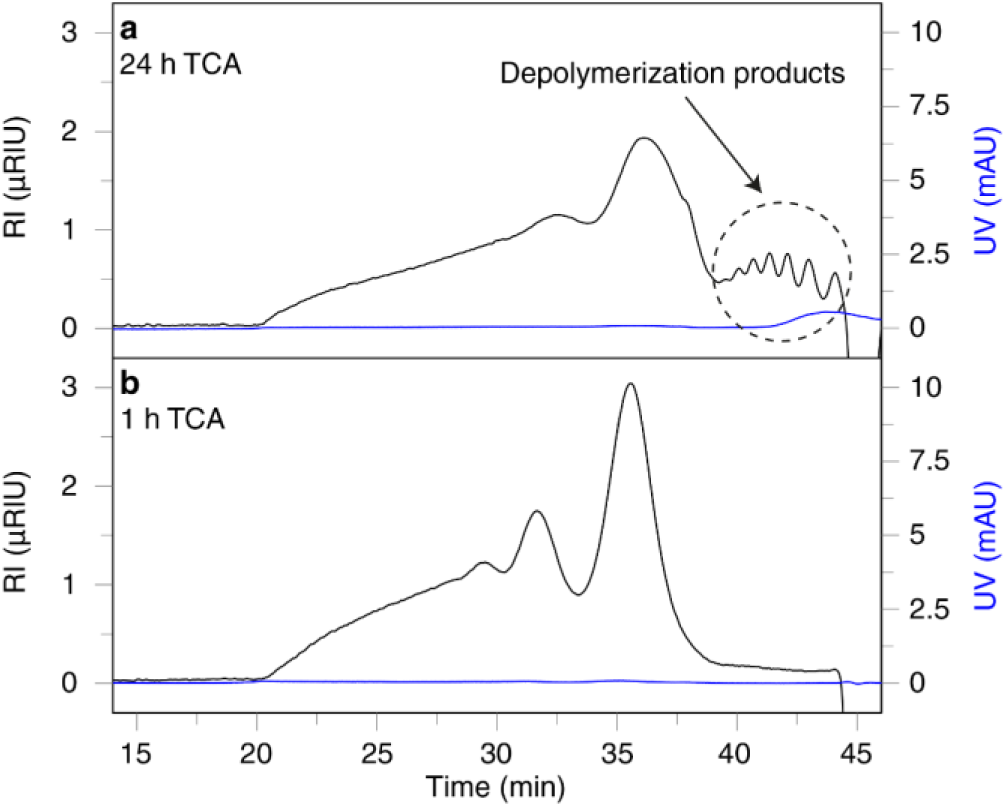
Observed CWPS hydrolysis by prolonged exposure to TCA. Size-exclusion chromatography (SEC) profiles with refractive index detection (RI, black traces) and UV detection (mAU_280nm_, blue traces) of CWPS isolates from *L. cremoris* TF121 exposed to 5% TCA for (a) 24 h and (b) 1 h at 4 °C during workup.

**Extended Figure 3.**
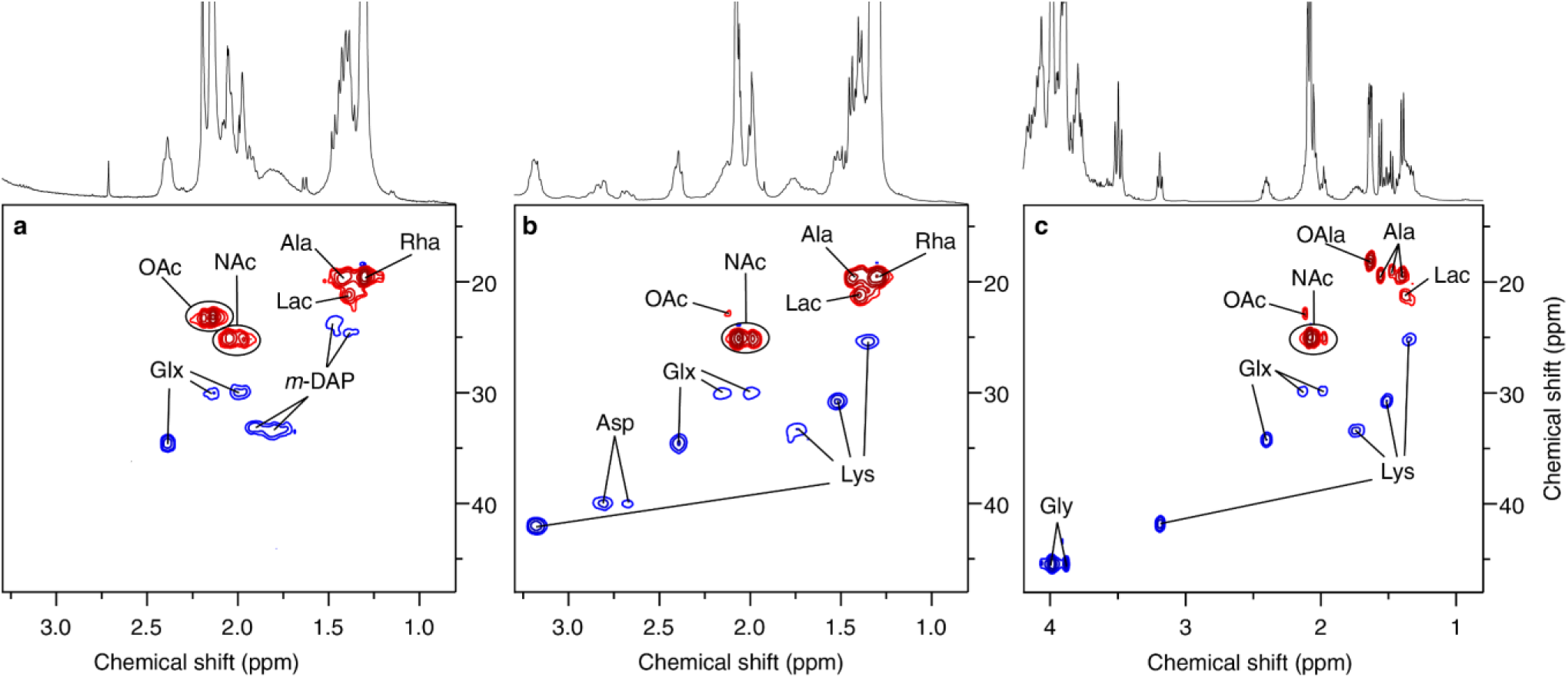
Identification of the PG residues by NMR. The figure shows multiplicity-edited HSQC spectra depicting the high-field region of **(a)** the high M_p_ CWPS from *L. plantarum* WCFS1 **(b)** CWG from *L. cremoris* TF121, and **(c)** *S. aureus* Newman. Signals stemming from CH_3_ groups are depicted in red, while signals from CH_2_ groups are blue. Glx: Isoglutamyl or γ-glutamyl, OAc: *O*-Acetyl, NAc: *N*-Acetyl, *m-*DAP: *meso*-Diaminopimeloyl, Rha: Rhamnosyl, Lys: Lysyl, Lac: Lactyl, Ala: Alanyl, OAla: *O*-Alanyl, Gly: Glycyl.

**Extended Figure 4:**
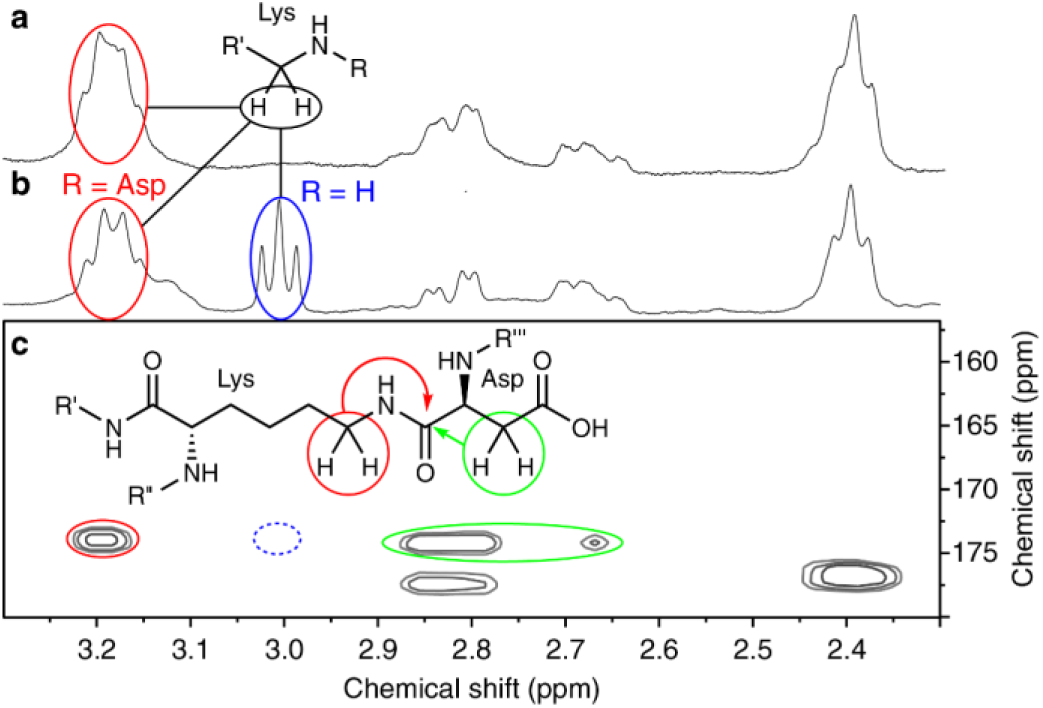
Identification of the PG crosslinking by NMR. **(a)** ^1^H NMR spectrum depicting the high-field region of the CWPS from *L. cremoris* TF121. **(b)** ^1^H NMR spectrum depicting the high-field region of the CWPS from *L. cremoris* CF117. **(c)** Excerpt from the HMBC spectrum of *L. cremoris* CF117 CWPS showing the Lys-Asp linkage for the signal at 3.19 ppm, which is not present for the signal at 3.00 ppm.

**Extended Figure 5.**
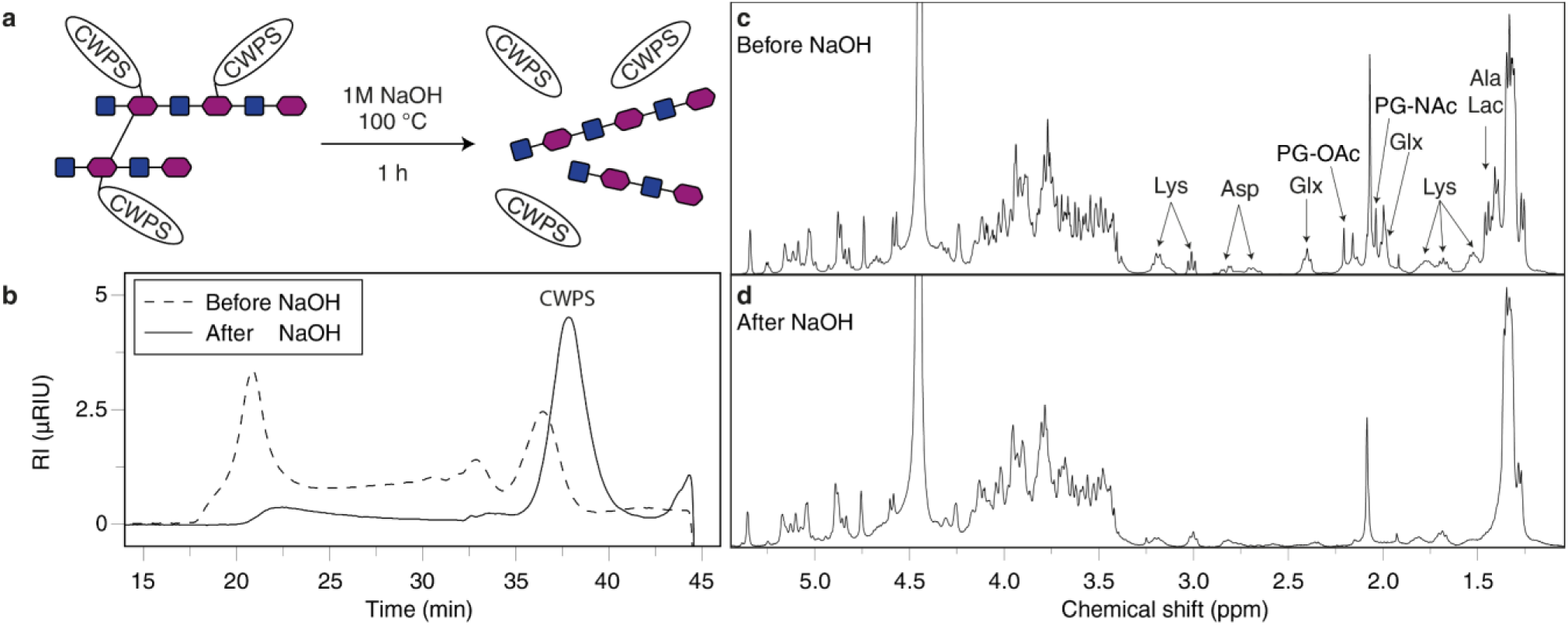
Removal of PG residues from *L. cremoris* CF117 CWPS by alkaline conditions. **(a)** Schematic representation of the reaction and conditions. Filled blue squares connected to filled magenta hexagons illustrate short fragments of the PG chain. Oval labelled ‘CWPS’ illustrate the CWPS polymer. **(b)** Size-exclusion chromatography (SEC) profiles of the CWPS extract with refractive index detection before and after alkaline treatment. **(c)** ^1^H NMR spectrum (400 MHz, D_2_O, 333 K) of the CWPS extract before alkaline treatment. **(d)** ^1^H NMR spectrum (400 MHz, D_2_O, 333 K) of the CWPS extract after alkaline treatment. Glx: Isoglutamyl or γ-Glutamyl, Lac: lactyl, Ala: Alanyl.

**Extended Table 1:**
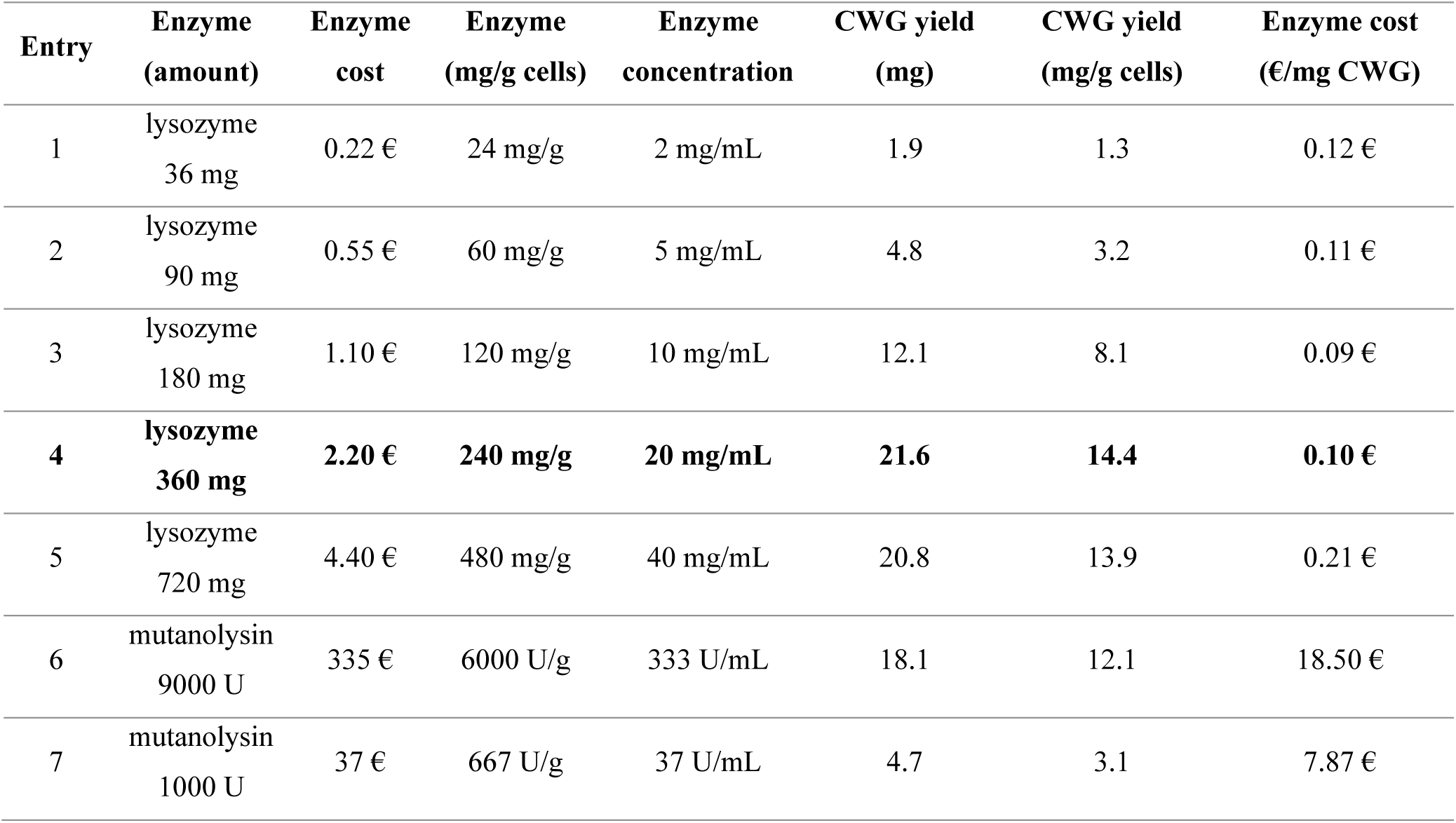
Optimization of enzymatic CWG extraction from *Lactococcus lactis* TF121. All extractions were performed on 1.5 g wet cells suspended in 18 mL water over 16 h at 37 °C. The enzyme cost is based on the prices for sigma- aldrich products 62971-50G-F (Lysozyme)^29^ and M9901-10KU (Mutanolysin)^30^ and does not include shipping or taxes (Values were retrieved on 24.09.2026).

## Notes

### Competing Interest Statement

Gordon Jacob Boehlich, Ahmad Tsjokajev, Pascal Mrozek, Gustav Vaaje-Kolstad and Bjorge Westereng have submitted a patent application covering the CWG extraction method described in this study. For the purpose of open access, the author has applied a CC BY public copyright license to any Author Accepted Manuscript (AAM) version arising from this submission.

## References

1. Gao, S. et al. Bacterial capsules: Occurrence, mechanism, and function. npj Biofilms Microbiomes 10, 21 (2024).

2. Khadka, S., Kinney, E. L., Ryan, B. E. & Mike, L. A. Mechanisms governing bacterial capsular polysaccharide attachment and chain length. Annals of the New York Academy of Sciences 1548, 80–98 (2025).

3. Guérin, H., Kulakauskas, S. & Chapot-Chartier, M.-P. Structural variations and roles of rhamnose-rich cell wall polysaccharides in Gram-positive bacteria. Journal of Biological Chemistry 298, 102488 (2022).

4. Rajagopal, M. & Walker, S. Envelope Structures of Gram-Positive Bacteria. in Protein and Sugar Export and Assembly in Gram-positive Bacteria (eds Bagnoli, F. & Rappuoli, R.) 1–44 (Springer International Publishing, Cham, 2017). doi:10.1007/82_2015_5021.

5. Larson, T. R. & Yother, J. Streptococcus pneumoniae capsular polysaccharide is linked to peptidoglycan via a direct glycosidic bond to β-D-N-acetylglucosamine. Proceedings of the National Academy of Sciences 114, 5695–5700 (2017).

6. Zeidan, A. A. et al. Polysaccharide production by lactic acid bacteria: from genes to industrial applications. FEMS Microbiol Rev 41, S168–S200 (2017).

7. Poxton, I. R. Chapter 5 - Teichoic Acids, Lipoteichoic Acids and Other Secondary Cell Wall and Membrane Polysaccharides of Gram-Positive Bacteria. in Molecular Medical Microbiology (Second Edition) (eds Tang, Y.- W., Sussman, M., Liu, D., Poxton, I. & Schwartzman, J.) 91–103 (Academic Press, Boston, 2015). doi:10.1016/B978-0-12-397169-2.00005-6.

8. Weidenmaier, C. & Peschel, A. Teichoic acids and related cell-wall glycopolymers in Gram-positive physiology and host interactions. Nat Rev Microbiol 6, 276–287 (2008).

9. Mistou, M.-Y., Sutcliffe, I. C. & van Sorge, N. M. Bacterial glycobiology: rhamnose-containing cell wall polysaccharides in Gram-positive bacteria. FEMS Microbiol Rev 40, 464–479 (2016).

10. Shen, Y. et al. Structural and functional diversity in Listeria cell wall teichoic acids. Journal of Biological Chemistry 292, 17832–17844 (2017).

11. Brown, S., Santa Maria, J. P. & Walker, S. Wall Teichoic Acids of Gram-Positive Bacteria. Annual Review of Microbiology 67, 313–336 (2013).

12. Pasquina, L. W., Santa Maria, J. P. & Walker, S. Teichoic acid biosynthesis as an antibiotic target. Current Opinion in Microbiology 16, 531–537 (2013).

13. Wang, H. et al. Discovery of Wall Teichoic Acid Inhibitors as Potential Anti-MRSA β-Lactam Combination Agents. Chemistry & Biology 20, 272–284 (2013).

14. Burns, K., Dorfmueller, H. C., Wren, B. W., Mawas, F. & Shaw, H. A. Progress towards a glycoconjugate vaccine against Group A Streptococcus. npj Vaccines 8, 48 (2023).

15. van Sorge, N. M. et al. The Classical Lancefield Antigen of Group A Streptococcus Is a Virulence Determinant with Implications for Vaccine Design. Cell Host & Microbe 15, 729–740 (2014).

16. Zorzoli, A. et al. Group A, B, C, and G Streptococcus Lancefield antigen biosynthesis is initiated by a conserved α-d-GlcNAc-β-1,4-l-rhamnosyltransferase. Journal of Biological Chemistry 294, 15237–15256 (2019).

17. Dalen, R. van, Peschel, A. & Sorge, N. M. van. Wall Teichoic Acid in Staphylococcus aureus Host Interaction. Trends in Microbiology 28, 985–998 (2020).

18. Sadovskaya, I. et al. Another Brick in the Wall: a Rhamnan Polysaccharide Trapped inside Peptidoglycan of Lactococcus lactis. mBio 8, 10.1128/mbio.01303-17 (2017).

19. Parlindungan, E. et al. Novel cell wall polysaccharide genotypes and structures of lactococcal strains isolated from milk and fermented foods. International Journal of Food Microbiology 424, 110840 (2024).

20. Chapot-Chartier, M.-P. et al. Cell Surface of *Lactococcus lactis* Is Covered by a Protective Polysaccharide Pellicle*. Journal of Biological Chemistry 285, 10464–10471 (2010).

21. Mahony, J. et al. The CWPS Rubik’s cube: Linking diversity of cell wall polysaccharide structures with the encoded biosynthetic machinery of selected Lactococcus lactis strains. Molecular Microbiology 114, 582–596 (2020).

22. Michon, F. & Uitz, C. Method for purifying polysaccharides. (2013).

23. Sadovskaya, I. & Guérardel, Y. Simple Protocol to Purify Cell Wall Polysaccharide from Gram-Positive Bacteria and Assess Its Structural Integrity. in Bacterial Polysaccharides: Methods and Protocols (ed. Brockhausen, I.) 37–45 (Springer, New York, NY, 2019). doi:10.1007/978-1-4939-9154-9_4.

24. Choudhury, B. et al. The Structure of the Major Cell Wall Polysaccharide of *Bacillus anthracis* Is Species- specific*. Journal of Biological Chemistry 281, 27932–27941 (2006).

25. Edgar, R. J. et al. Discovery of glycerol phosphate modification on streptococcal rhamnose polysaccharides. Nat Chem Biol 15, 463–471 (2019).

26. Guerardel, Y. et al. Complete Structure of the Enterococcal Polysaccharide Antigen (EPA) of Vancomycin-Resistant Enterococcus faecalis V583 Reveals that EPA Decorations Are Teichoic Acids Covalently Linked to a Rhamnopolysaccharide Backbone. mBio 11, 10.1128/mbio.00277-20 (2020).

27. Yokogawa, K., Kawata, S., Nishimura, S., Ikeda, Y. & Yoshimura, Y. Mutanolysin, Bacteriolytic Agent for Cariogenic Streptococci: Partial Purification and Properties. Antimicrobial Agents and Chemotherapy 6, 156– 165 (1974).

28. Ghosh, R. & Cui, Z. F. Purification of lysozyme using ultrafiltration. Biotechnology and Bioengineering 68, 191– 203 (2000).

29. Lysozym aus Hühnereiweiss powder (crystalline), 70000-140000 U/mg | Sigma-Aldrich. https://www.sigmaaldrich.com/DE/de/product/sial/62971.

30. Mutanolysin aus Streptomyces globisporus ATCC 21553 lyophilized powder, ≥4000 units/mg protein (biuret), Chromatographically purified | Sigma-Aldrich. https://www.sigmaaldrich.com/DE/de/product/sigma/m9901.

31. Vinogradov, E. et al. Structural studies of the cell wall polysaccharide from *Lactococcus lactis* UC509.9. Carbohydrate Research 461, 25–31 (2018).

32. Ainsworth, S. et al. Differences in Lactococcal Cell Wall Polysaccharide Structure Are Major Determining Factors in Bacteriophage Sensitivity. mBio 5, 10.1128/mbio.00880-14 (2014).

33. Vinogradov, E. et al. Determination of the cell wall polysaccharide and teichoic acid structures from *Lactococcus lactis* IL1403. Carbohydrate Research 462, 39–44 (2018).

34. Ajay Castro, S., et al. Recombinant production platform for Group A Streptococcus glycoconjugate vaccines. npj Vaccines 10, 16 (2025).

35. Kessler, R. E., van de Rijn, I. & McCarty, M. Characterization and localization of the enzymatic deacylation of lipoteichoic acid in group A streptococci. J Exp Med 150, 1498–1509 (1979).

36. Jansson, P.-E., Lindberg, B., Andersson, M., Lindquist, U. & Henrichsen, J. Structural studies of the capsular polysaccharide from *streptococcus pneumoniae* type 2, a reinvestigation. Carbohydrate Research 182, 111–117 (1988).

37. Gaikwad, W. K., Jana, S. K., Dhere, R. M., Ravenscroft, N. & Kodam, K. M. Partial depolymerization of capsular polysaccharides isolated from *Streptococcus pneumoniae* serotype 2 by various methods. Carbohydrate Research 512, 108503 (2022).

38. Gisch, N. et al. Structural Reevaluation of Streptococcus pneumoniae Lipoteichoic Acid and New Insights into Its Immunostimulatory Potency *. Journal of Biological Chemistry 288, 15654–15667 (2013).

39. Johnson, S. D., Lacher, K. P. & Anderson, J. S. Carbon-13 NMR spectroscopic study of teichuronic acid from Micrococcus luteus cell walls. Comparison of the polysaccharide isolated from cells with that synthesized in vitro. Biochemistry 20, 4781–4785 (1981).

40. Hase, S. & Matsushima, Y. Structural Studies on a Glucose-containing Polysaccharide Obtained from Cell Walls of Micrococcus lysodeikticus. The Journal of Biochemistry 72, 1117–1128 (1972).

41. Shashkov, A. S., Potekhina, N. V., Senchenkova, S. N. & Kudryashova, E. B. Anionic polymers of the cell wall of Bacillus subtilis subsp. subtilis VKM B-501T. Biochemistry Moscow 74, 543–548 (2009).

42. Mistretta, N. et al. Glycosylation of Staphylococcus aureus cell wall teichoic acid is influenced by environmental conditions. Sci Rep 9, 3212 (2019).

43. Tomita, S., Tanaka, N. & Okada, S. A rapid NMR-based method for discrimination of strain-specific cell wall teichoic acid structures reveals a third backbone type in Lactobacillus plantarum. FEMS Microbiol Lett 364, fnx034 (2017).

44. Wiull, K. et al. Antigen surface display in two novel whole genome sequenced food grade strains, Lactiplantibacillus pentosus KW1 and KW2. Microb Cell Fact 23, 19 (2024).

45. Schleifer, K. H. & Kandler, O. Peptidoglycan types of bacterial cell walls and their taxonomic implications. Bacteriological Reviews 36, 407–477 (1972).

46. Bera, A., Herbert, S., Jakob, A., Vollmer, W. & Götz, F. Why are pathogenic staphylococci so lysozyme resistant? The peptidoglycan O-acetyltransferase OatA is the major determinant for lysozyme resistance of Staphylococcus aureus. Molecular Microbiology 55, 778–787 (2005).

47. Vidal, M.-L., Gautron, J. & Nys, Y. Development of an ELISA for Quantifying Lysozyme in Hen Egg White. J. Agric. Food Chem. 53, 2379–2385 (2005).

48. Tsjokajev, A. et al. Pilot-scale production, purification, and structural characterization of a glycosaminogalactan exopolysaccharide from *Lactiplantibacillus pentosus*. Carbohydrate Polymers 381, 125125 (2026).

49. Tsjokajev, A., Boehlich, G. J., Horn, S. J., Vaaje-Kolstad, G. & Westereng, B. High molecular-weight polysaccharide contamination from yeast extract in semi-defined bacteriological media: Effects on exopolysaccharide production and purity. 2026.02.27.708493 Preprint at 10.64898/2026.02.27.708493 (2026).

50. Vander Heyden, Y., Popovici, S. T. & Schoenmakers, P. J. Evaluation of size-exclusion chromatography and size- exclusion electrochromatography calibration curves. Journal of Chromatography A 957, 127–137 (2002).

51. Fontana, C. & Widmalm, G. Primary Structure of Glycans by NMR Spectroscopy. Chem. Rev. 123, 1040–1102 (2023).

52. Speciale, I. et al. Liquid-state NMR spectroscopy for complex carbohydrate structural analysis: A hitchhiker’s guide. Carbohydrate Polymers 277, 118885 (2022).

